# Regulation of modulatory cell activity across olfactory structures in *Drosophila melanogaster*

**DOI:** 10.1101/522177

**Authors:** Xiaonan Zhang, Kaylynn Coates, Andrew Dacks, Cengiz Gunay, J. Scott Lauritzen, Feng Li, Steven A. Calle-Schuller, Davi Bock, Quentin Gaudry

## Abstract

All centralized nervous systems possess modulatory neurons that arborize broadly across multiple brain regions. Such modulatory systems are critical for proper sensory, motor, and cognitive processing. How single modulatory neurons integrate into circuits within their target destination remains largely unexplored due to difficulties in both labeling individual cells and imaging across distal parts of the CNS. Here, we take advantage of an identified modulatory neuron in *Drosophila* that arborizes in multiple olfactory neuropils. We demonstrate that this serotonergic neuron has opposing odor responses in its neurites of the antennal lobe and lateral horn, first and second order olfactory neuropils respectively. Specifically, processes of this neuron in the antennal lobe have responses that are inhibitory and odor-independent, while lateral horn responses are excitatory and odor-specific. The results show that widespread modulatory neurons may not function purely as integrate-and-fire cells, but rather their transmitter release is locally regulated based on neuropil. As nearly all vertebrate and invertebrate neurons are subject to synaptic inputs along their dendro-axonic axis, it is likely that our findings generalize across phylogeny and other broadly-projecting modulatory systems.

**Significance:** The centrifugal innervation of neuronal circuits is ubiquitous across centralized nervous systems. Such inputs often arise from modulatory neurons that arborize broadly throughout the brain. How information is integrated in such cells and how release from their distant terminals is regulated remains largely unknown. We show that a serotonergic neuron that innervates multiple stages of odor processing in *Drosophila* has distinct activity throughout its neurites, including opposite polarity responses in first and second order olfactory neuropils. Disparate activity arises from local interactions within each target region. Our results show that such neurons exhibit dendritic computation rather than somatic integration alone, and that examining local interactions at release sites is critical for understanding centrifugal innervation.

## Introduction

Virtually all neuronal circuits are subject to neuromodulation from both neurons intrinsic to a network and extrinsic centrifugal sources (1, 2). In vertebrates, extrinsic modulation is often supplied by nuclei located deep within the brainstem that release a variety of transmitters such as norepinephrine (NE) (3), serotonin (5-HT) (4, 5), dopamine (DA) (6, 7), or acetylcholine (Ach) (8). For example, the olfactory bulb (OB) in mammals receives a tremendous amount of centrifugal innervation (9) that can be critical for proper olfactory behavior (10). However, by spanning and innervating most cortical and subcortical regions, modulatory systems target multiple points along the sensory-motor axis of functional circuits. A prominent view of such modulatory systems is that they provide a mechanism for top-down regulation of sensory processing (11, 12) and that they help coordinate activity across brain regions (13). Modulatory systems are traditionally regarded as integrate-and-fire models where the neurons integrate synaptic inputs in their dendrites within their local nuclei and use action potentials to broadcast this signal to release sites. Such models imply that modulator release will be inherently correlated across distal targets within the brain. However, as virtually all axons are subject to pre-synaptic regulation (14), it is likely that most centrifugal modulatory neurons are subject to local influences by the circuits that they infiltrate. This implies that the local release of transmitters from such systems may instead be decorrelated across brain regions. Decorrelating transmitter release across brain regions is advantageous, as it would provide greater flexibility in how neuromodulation may be employed. How transmitter release is locally regulated in modulatory neurons and how signals propagate through their processes has been exceedingly difficult to study in vertebrate systems due to many contributing factors. First, the vertebrate cerebrum is large and imaging the extensive processes of such neurons across brain areas requires specialized tools (15–18). Second, individual modulatory neurons within the same brainstem nucleus are highly heterogeneous in their projection patterns (19, 20) making it difficult to assign activity across brain regions to individual cells. Additionally, the spatial pattern of extrinsic input activation can also be stimulus specific. For example, different odors can activate unique presynaptic terminals from piriform neurons that feed back into the OB (21). But whether such odor responses in centrifugal inputs arise from the recruitment of different individual cortical neurons or from local axo-axonic interactions within the OB remains unknown. The spatial pattern of cholinergic into the OB is also odor-specific and arises through similarly undescribed mechanisms (22).

The *Drosophila* brain is an ideal preparation to study how signals propagate through wide-field modulatory neurons because such cells are often stereotyped and can be genetically targeted across individual flies. For example, only one serotonergic neuron per hemisphere, termed the contralaterally-projecting serotonin-immunoreactive deuterocerebral neuron (CSDn), innervates both the first and second order olfactory neuropils in the fly brain (23–26) (Fig. 1*A*). CSDns can be targeted genetically (26) and are critical for various olfactory behaviors involving appetitive (27) and pheromonal odorants, including cis-vaccenyl acetate (cVA) (28). The modulation of cVA-evoked behavioral responses is especially interesting because the CSDns are thought to participate in top-down modulation and to have their effects mainly in the first olfactory relay, the antennal lobe (AL) (24, 28, 29). However, CSDn processes avoid the cVA-sensitive DA1 glomerulus (25, 30) and DA1 projection neuron (PN) odor responses are not modulated with strong CSDn stimulation (30). Finally, whole-cell recordings show strong inhibition of the CSDn during stimulation with cVA (30). This suggests that the modulatory effects of the CSDns on cVA-guided behavior may not occur in the AL, but rather in another olfactory neuropil that the CSDns innervate. This suggests that the modulatory effects of the CSDns on cVA-guided behavior may not occur in the AL, but rather in one of the other olfactory neuropil that the CSDns innervate. Because the CSDns express pre-and postsynaptic markers throughout their arborizations (30), it is possible that transmitter release is locally regulated via inputs from their target networks (31) and that olfactory-mediated modulation occurs predominantly downstream of the AL. In this study we employed 2-photon calcium imaging, electron microscopy, and compartmental modeling to show that the CSDns integrate synaptic inputs locally within their target regions giving rise to distinct odor evoked activity patterns within different olfactory neuropil.

**Figure 1.**
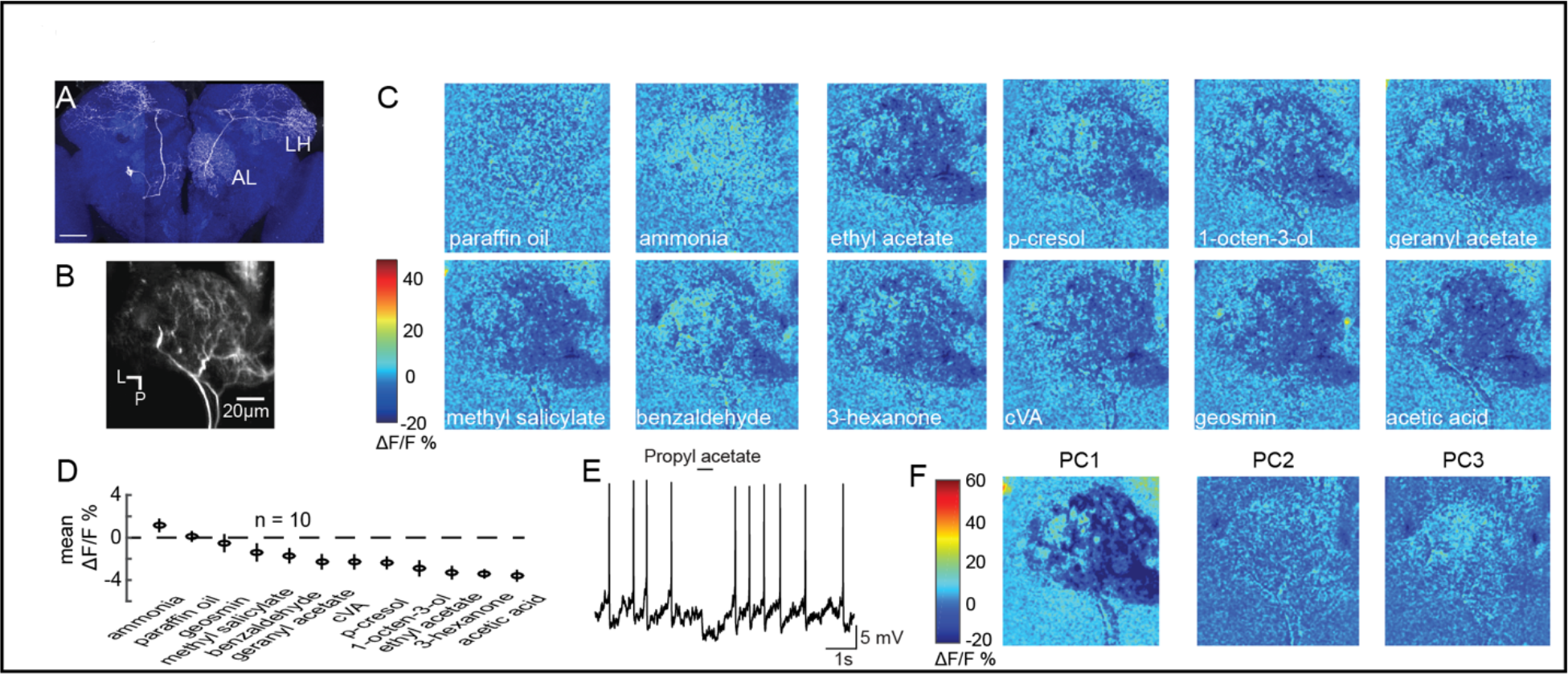
Olfactory stimulation inhibits CSDn processes in the AL. (*A*) (*A*) A single CSDn expressing GFP shows processes in both the AL and the LH. White = GFP expression, blue = neuropil labeling via N-cadherin immunocytochemistry. Scale bar = 50um. (*B*) Background fluorescence from GCaMP6s expression in CSDn neurites in the AL. L = lateral, P = posterior. The same notation is used in subsequent figures (*C*) Odor-evoked changes in calcium (ΔF/F) levels in the AL processes of the CSDn. Cooler colors (blue) represent decreases in calcium levels and warmer (red) colors show increases in calcium. Odors are diluted 10^−2^ in paraffin oil (PA), which responses serve as a solvent control. Images are scaled and oriented as in *B*. (*D*) Odor responses ranked according to the strength of the observed inhibition (n = 10). (*E*) Odor-evoked inhibition observed in the CSDn soma via whole-cell patch-clamp recording. (*F*) Principal component analysis performed on the spatial pattern at the peak of the odor responses. The first three PCs are shown. A structured response is only observed in PC1. Images are scaled and oriented as in *B*.

## Results

To examine how CSDns contribute to olfactory processing across brain regions, we employed GCaMP6s (32) and 2-photon volumetric microscopy to characterize olfactory responses across their arbors. We initially imaged CSDn neurites in the AL (Fig. 1*B*) and found that nearly all compounds in a diverse panel of odorants resulted in inhibition (Fig. 1*C* and *D*). The only exception was ammonia, which produced a weak level of excitation as previously reported in whole-cell somatic recordings (30). These results are consistent with previous studies demonstrating that odors generally inhibit the CSDns during electrophysiological somatic recordings (Fig. 1*E*) (30), which is likely due to prominent input from GABAergic local interneurons (LN) (25, 30).

Inhibition in the AL scales with increasing odor intensity and the spatial pattern of activation of GABAergic LNs is odor invariant (33). Inhibition of the CSDn also scales with odor intensity (30) but it is unknown if unique odors can recruit distinct spatial patterns of CSDn activity. We used principal components analysis (PCA) to examine the spatial profile of CSDn inhibition in response to our odor panel. This analysis revealed that only the first PC generated an structured image showing inhibition in the AL while PC2 captured the excitation resulting from stimulation with ammonia (Fig. 1*F*). Thus, aside from ammonia, CSDn processes in the AL receive odor-invariant inhibition, and this inhibition scales with odor intensity (30).

We next examined CSDn processes in the lateral horn (LH; Fig. 2*A*), a brain region that receives direct input from the AL and mediates innate olfactory behaviors (34). Surprisingly, we found that every odorant in our panel produced excitation in the CSDn LH arbors (Fig. 2*B*). Furthermore, PCA revealed that CSDn odor responses in the LH varied spatially (Fig. 2*C*) and displayed a greater coefficient of variation compared to responses in the AL (Fig. 2*D*). These results show that the processes of the CSDn have opposing responses to odor stimulation across different olfactory regions (Fig. 2*F*). The CSDn has both pre-and postsynaptic sites in the LH (30), so GCaMP signaling could represent either the activation of CSDn postsynaptic receptors or calcium influx at presynaptic release sites. Some GCaMP signaling in the LH must represent local activation of postsynaptic receptors in the CSDn since its AL processes are simultaneously inhibited. To assess whether this activity also correlates with synaptic release from the CSDn, we employed sytGCaMP6s, a variant of the calcium sensor that is tethered to synaptotagmin and trafficked to presynaptic release sites (35). Olfactory stimulation showed increased sytGCaMP6s signaling (Fig. S*1*), suggesting odorants likely evoke transmitter release in the LH processes of the CSDn.

**Figure 2.**
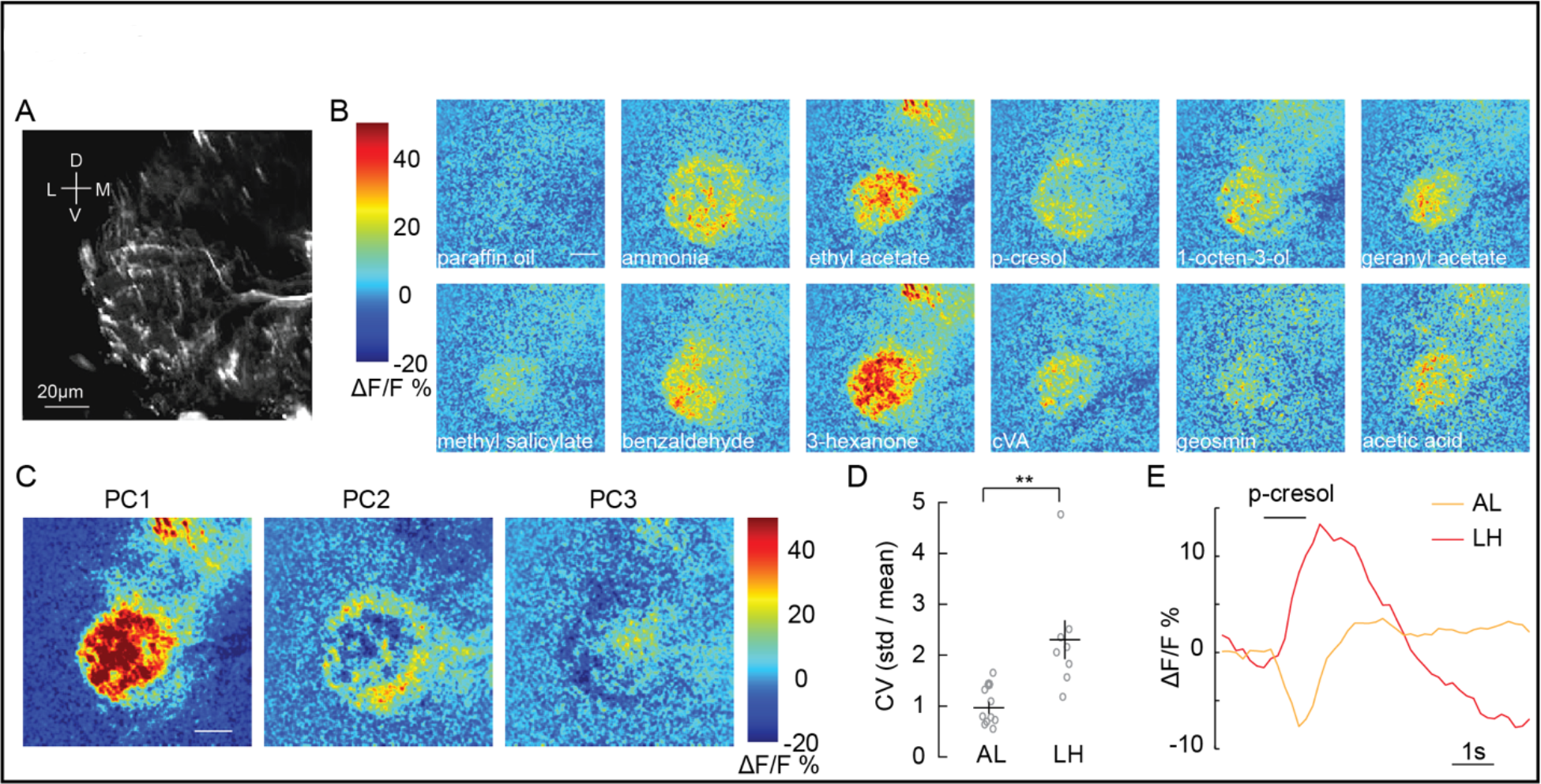
Olfactory stimulation excites CSDn processes in the LH. (*A*) Background fluorescence from GCaMP6s expression in CSDn neurites in the LH. (*B*) Odor-evoked changes in calcium observed in CSDn processes in the LH. Odor scale bar = 20 μm (*C*) Principal component analysis performed on the spatial pattern at the peak of the odor responses for a sample preparation. The first three PCs are shown. A structured response is observed in each PC. Images are scaled as in *B*. (*D*) The coefficient of variation for odor responses in the AL and LH. n = 10, AL and n = 8, LH. p < 0.01, Student’s t-test. (*E*) A comparison of the time series of ΔF/F responses in the AL and LH to a single odorant.

To determine if spatial patterns of CSDn odor-evoked activity in the LH were odor-specific, we quantified the similarity between CSDn odor responses by first computing the spatial correlation between them (Fig. 3*A*). We then calculated the Euclidean linkage distance between all odor correlations to illustrate which odors are most similarly encoded in the neurites of the CSDn in the LH (Fig. 3*B*). How might odor specific responses arise in the CSDn processes in the LH? PNs are a potential source of excitatory input to the CSDns in the LH as they are cholinergic, and their axons segregate anatomically (36, 37) and functionally (38–40) in this region. We compared the odor representations of PN and CSDn processes within this structure to determine if PNs may provide excitatory drive to the CSDn branches in the LH. The linkage distance between the spatial pattern of odor responses was highly correlated between PN and CSDn responses in the LH (Fig. S*2* and 3*C*). These data suggest that PNs may provide direct synaptic input onto the CSDn locally in the LH. Using synaptobrevin GFP Reconstitution Across Synaptic Partners (syb:GRASP) (41, 42) we observed a positive signal for PNs synapsing onto the CSDns (Fig.3*D-F*) in the LH. We further verified this synaptic connection by looking at the connectivity of a CSDn reconstructed within a whole fly brain EM dataset (43). PNs from at least 17 glomeruli (presynaptic PN tracings provided as a personal communication from Greg Jefferis, Phillip Schlegel, Alex Bates, Marta Costa, Fiona Love and Ruari Roberts) show direct input to the CSDn throughout the LH (Fig. 3 *G-I*). Together these results suggest that the CSDns receives input from different cell classes in the AL and LH to shape its olfactory responses locally. In the AL, GABAergic LNs constitute a major input to the CSDns and drive inhibitory responses independent of odor identify. In the LH, olfactory responses in the CSDn neurites are excitatory, odor specific, and are likely in part by direct PN input.

**Figure 3.**
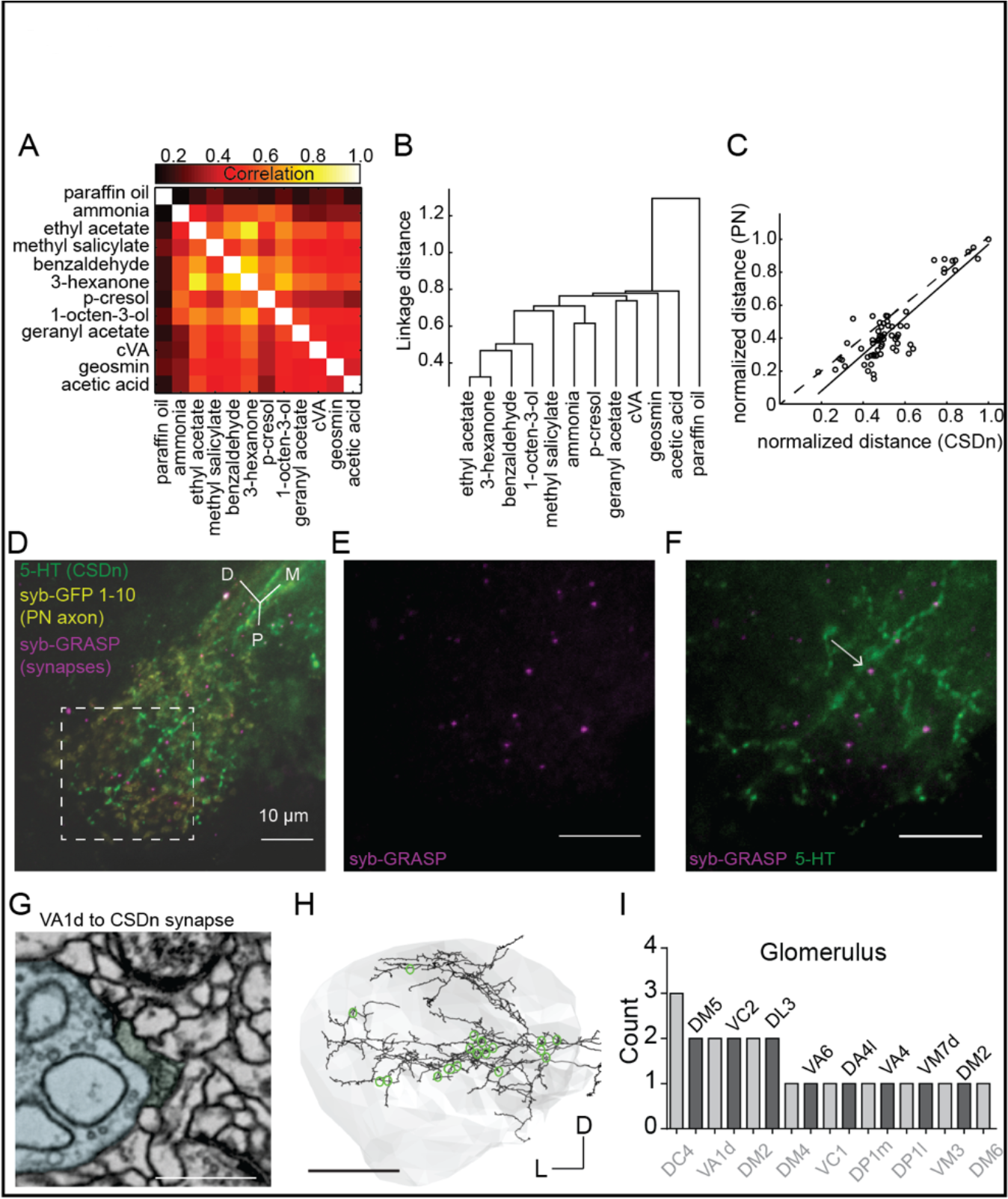
Projection neurons shape CSDn responses in the LH via direct synaptic input. (*A*) A cross correlation of the spatial profile of odor responses in CSDn LH processes. (*B*) Clustering analysis using the Euclidean distance between correlations for odor pairs from (*A*). (*C*) Regression analysis on correlation distances between odor responses in processes of the CSDn and PN axons in the LH (R^2^= 0.73, p<0.001, n = 10 preparations for CSDn and n = 5 for PN responses). (*D – F*) GRASP images showing direct synaptic input from PNs onto CSDn terminals. Green = 5-HT antibody to label CSDn processes, yellow = syb-GFP subunits 1-10 expressed in PN axons, magenta = syb-GRASP labeling of synapses. Scale bars = 20 μm. (*G*) EM image of a direct VA1d PN synapse onto a CSDn neurite in the AL Scale bar = 500nm. (*H*) EM reconstruction of the CSDn (black) arbors in the LH (grey boundary). Location of individual PN synapses onto the CSDn are marked in green. Scale bar = 25um. (*I*) A total count of PN onto CSDn synapses in the LH separated by glomerular identity. Glomeruli are listed above and below bars for clarity. Counts taken from 1 female brain.

The function of a neuron is often dictated by the manner in which synaptic inputs are integrated across its dendritic arbor. We therefore asked whether the AL and LH neurites of the CSDn function as electrotonically independent compartments within the same cell, or if signals propagate between regions during odor sampling. To examine how voltage spreads throughout the CSDn, we built a passive compartmental model based on an anatomical reconstruction of the CSDn generated from the whole fly brain EM dataset (43, 44) (Fig. 4*A*, S*3*). The model was generated by adjusting the membrane capacitance (C_m_), the membrane conductance (g_leak_), and the axial resistance (R_a_) so that simulated current injections into the model soma matched physiological responses taken *in vivo* (Fig. 4*B*). A wide range of models with varying properties provided reasonable fits to the *in vivo* CSDn recordings. In these models, injecting simulated hyperpolarizing IPSPs into the AL resulted in varying amounts of spread throughout the neurites of the CSDn. In some models, the hyperpolarization was constrained only to the local injection site (Fig. 4*C*), while in other models the hyperpolarization spread throughout the AL (Fig. 4*D*). However, whole-cell recordings *in vivo* show that the inhibition arising from the AL indeed spreads to the soma (30). Several of the models that we generated displayed this property. Importantly, all models that displayed somatic inhibition when current was injected into the CSDn processes in the AL also showed inhibition in the LH (Fig. 4*E*). These results suggest that the geometry and passive properties of the CSDn allow the propagation of inhibition from the AL to the LH during olfactory stimulation.

**Figure 4.**
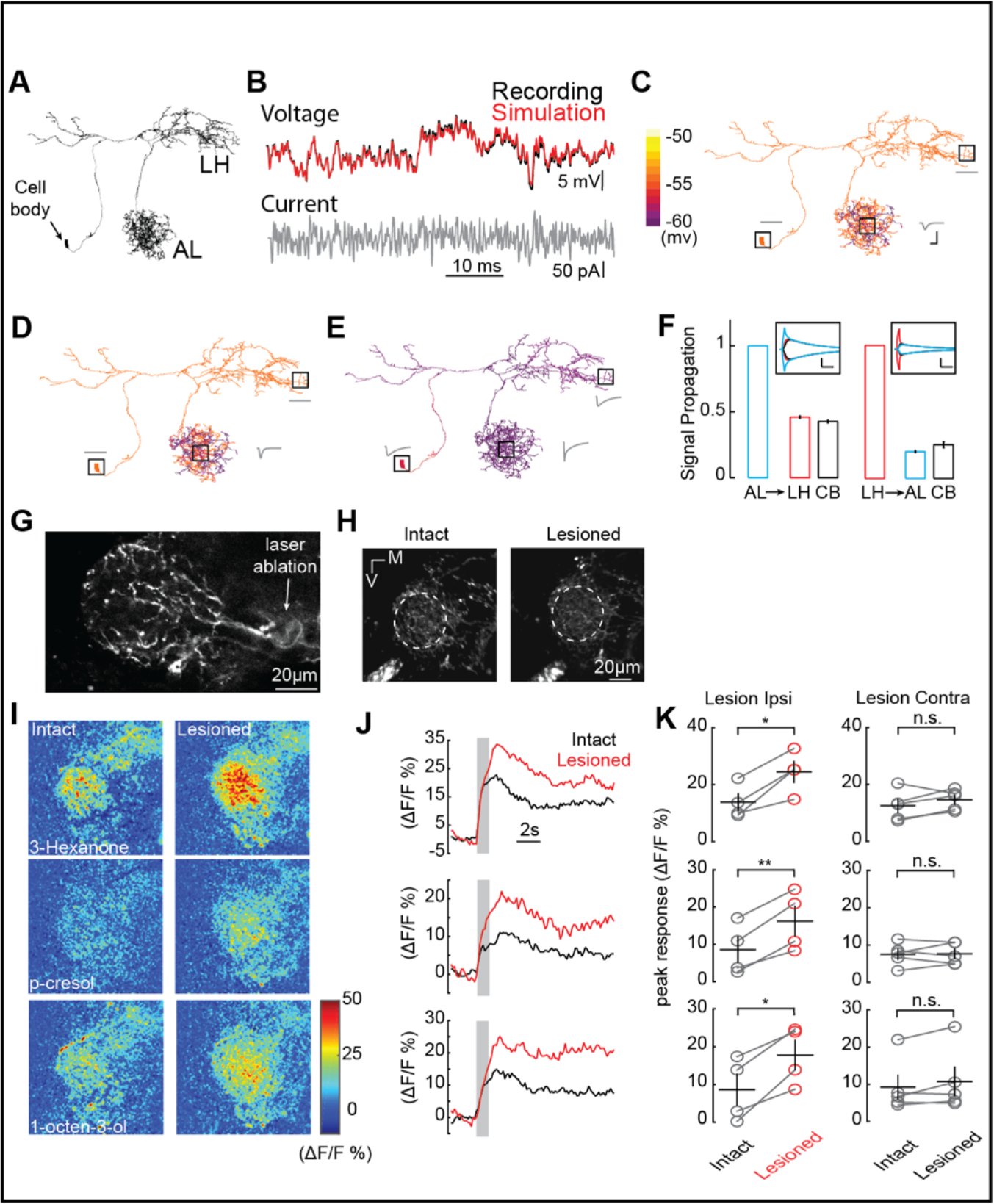
AL inhibition suppresses CSDn responses in the LH. (*A*) Morphological EM reconstruction of the CSDn arborizations used for a biophysical model consisting of 6056 compartments. (*B*) Somatic membrane potential changes in response to a white-noise current injection during an *in vivo* CSDn recording compared to responses of a passive compartmental model, to the same stimulus, fitted by optimizing three anatomical parameters, C_m_ (μF/cm^2^), g_leak_ (S/cm^2^), and R_a_ (Ωcm), (*C*) An example of a model with well-fit somatic responses where a hyperpolarizing current injection in the AL resulted in highly localized voltage changes in AL only. A time series of voltage responses are shown for the soma, AL, and LH as gray traces near those structures. C_m_ = 4.8,, g_leak_ = 3094.3, and R_a_ = 149.0183. Horizontal scale bar = 5 ms and vertical scale bar = 5 mV. (*D*) A similar model where injection of the current in the AL causes a voltage change throughout a greater portion of the AL. C_m_ = 4.8,, g_leak_ = 2911.3, and R_a_ = 28.83. Scale bars as in *C*. (*E*) A sample model where current injection into the AL causes both somatic and LH voltage changes. C_m_ = 1.8,, g_leak_ = 77.33, and R_a_ = 0.12. Scale bars as in *C*. (*F*) The proportion of voltage signal observed in the LH, AL, and cell body (CB) when voltage changes are induced in each region. The same model as in panel *E* was used to generate the data. *Left*, voltage steps are induced into the AL and voltage deflections are reported in the LH and CB. *Right*, voltage steps are induced in the LH and their effects are measured in the AL and CB. Horizontal scale bar = 10 ms and vertical scale bar = 10 mV.(*G*) A 2P image of basal GCaMP6s signal showing the effects of laser ablation in the CSDn LH neurite. (*H*) Basal GCaMP6s signals remain in the LH after laser ablation. (*I*) ΔF/F responses of CSDn neurites in the LH to a set of odorants before and after laser ablation. (*J*) A times series of the changes in calcium levels in response to odorants in (*I*). (*K*) *Left*, quantification of odor response amplitudes in CSDn LH neurites before and after laser ablation. *Right*, control responses when the contralateral processes of the CSDn where ablated. * = p < 0.05, ** = p < 0.01, n.s. = not significant and p > 0.05, paired Student’s t-tests.

The propagation of voltages through the dendrites of neurons is not always symmetrical and can be biased by impedance mismatches as well as differences in the diameters of intersecting branches (45). We therefore asked if voltage changes generated in the LH would propagate as efficiently to the AL compared to the opposite direction. We injected current into the either the LH or the AL of the model to elicit voltage changes of similar magnitudes and calculated the proportion of the signal that propagated to the other region. This approach revealed that the geometry of the CSDn arbors favor the propagation of voltage signals from the AL preferentially to the LH (Fig. 4*F*).

If voltage signals preferentially propagate from the AL to the LH, one would predict that isolating the CSDn processes in the LH from the rest of the CSDn would disinhibit any odor evoked excitation due to local input within the LH. We therefore used 2-photon laser ablation to sever the CSDn neurites just proximal to their entrance into the LH. This manipulation isolates the LH processes of the CSDn from the AL, while still allowing these neurites to respond to synaptic input within the LH (Fig. 4*G* and *H*). Olfactory responses to all odors tested increased in the processes of the CSDn in the LH following the removal of AL inhibition (Fig. 4*I*, *J*, and *K*). As a control for the non-specific damage of laser ablation, we severed the branches of the CSDn in the contralateral hemisphere and found that this had no impact on odor responses in the intact branches of the CSDn in the ipsilateral LH (Fig. 4*K*). These results demonstrate that during normal odor sampling, inhibition from the AL propagates to suppress olfactory responses in the LH.

## Discussion

There has been a recent interest in characterizing the structures that provide input to serotonergic neurons in vertebrates in an attempt to understand the types of processes that might impact 5-HT release (46). Such studies relied on genetically restricted retrograde labeling to identify regions that provide monosynaptic inputs to serotonergic raphe neurons (47–49). Our functional approach of calcium imaging across brain regions is complementary but has the advantage of revealing where signals are integrated and determining whether the net effect of that integration within is excitatory or inhibitory. Specifically, we have shown that a serotonergic neuron with broad arbors integrates locally at multiple neuropil along to the olfactory pathway in *Drosophila*. Interestingly, synaptic inputs at the first and second order processing stages of olfaction impose opposite polarity responses in this neuron and likely decorrelate synaptic release in distinct target regions. Inhibition dominates responses in the AL and odor-specific excitation is prominent in the downstream LH. This suggests that across a single widely projecting modulatory neuron, branches within distinct neuropils can operate in different manners.

### Local regulation of release in modulatory neurons

Classical methods of studying the activation of modulatory systems in vertebrates rely heavily on recording extracellular spikes in the nuclei that house the neurons’ somas. Such approaches promote a view of modulatory systems as comprising traditional integrate-and-fire neurons where excitatory and inhibitory inputs are assimilated only at the dendrites and conveyed to all release sites in a correlated manner. Our study demonstrates that the activity within a single serotonergic neuron can vary across neuropils involved in processing the same sensory modality. As the same modulator may perform distinct functions in different brain regions, local regulation of release allows these functions to be employed independently. For instance, 5-HT in the OB indirectly inhibits OSN terminals and has been proposed to serve a gain control function (50), while in the piriform cortex 5-HT has no effect on stimulus input, but rather only decreases spontaneous activity (51). Additionally, 5-HT has different effects on mitral cells in the main versus mitral cells in the accessory olfactory bulb (52). Local regulation of 5-HT release would allow these processes to be engaged in an independent and combinatorial manner, and thus allow for a greater net overall modulatory capacity.

Local regulation and decoupling of modulator release across synaptic sites is not unique to invertebrates and has been implicated in the normal function of the mammalian DA system. First, DA release is only partially correlated with firing activity, and release can be locally evoked in the absence of spiking in DA cells (53). Additionally, local inactivation of the nucleus accumbens (NA) decreases DA release in the NA without impacting DA neuron firing (54). This has led to the theory that DA release can signal both motivation and reward prediction errors (RPE) on similar timescales (55). Dopamine release related to motivation is thought to be shaped by local presynaptic mechanisms while dopamine related to RPE correlates more strongly with the firing properties of DA neurons (56). Measuring serotonergic transmission across release sites is more difficult compared to dopamine (57), nevertheless, it is highly likely that 5-HT is also regulated by local presynaptic mechanisms (58–61). Whether there is local regulation of 5-HT release in the vertebrate olfactory system is more controversial. EM reconstructions of raphe terminals in the OB have failed to reveal postsynaptic densities in raphe axons (62, 63). This may be because such input is extra synaptic (31), as is the case with GABAB receptors in the raphe nucleus proper (64).

### Multi-dendritic processing

Local integration in the AL and LH allow the CSDn to independently process and shape sensory information at multiple points in the early olfactory pathway of the fly. Specifically, we found that PN axons directly excite CSDn terminals in the LH in an odor specific manner while CSDn branches in the AL are inhibited. Previous studies have shown that the CSDn soma is also inhibited by odors and that stimulation of the CSDn has little impact on olfactory circuitry in the AL (30), despite the CSDn being critical for normal olfactory behavior (27, 28). Our current study resolves this issue by suggesting instead that the CSDn modulates behavior by affecting odor-processing in the LH (Fig. 4 and S*4*), which has previously been unexplored. What then is the purpose of odor-evoked inhibition in the AL? Our compartmental modeling shows that AL and LH processes of the CSDn are electrotonically connected, but that voltage preferentially passes from the AL to the LH. CSDn inhibition scales with increasing odor strength due to robust GABAergic LN input in the AL (25, 30), and this inhibition shunts LH responses proportionally. This configuration ultimately allows olfactory modulation to be odor specific while being less dependent on odor concentration. Multi-dendritic computing is critical for processing other sensory modalities as well (65), but most notably vision (66, 67). Synaptic integration between dendrites is used to compute object motion across the visual field on a collision course with the observer. Interestingly, the dendrites of starburst amacrine cells in the mammalian retina express mGluR to isolate dendritic compartments thus preventing integration to non-preferred stimuli while enabling regulated integration specifically to preferred directions of motion (67). Our laser ablation experiments demonstrate that CSDn neurites influence one another during olfaction, but it is intriguing that such coupling could be state dependent and regulated by active conductances.

### Top-down versus bottom-up neuromodulation

The CSDn was originally proposed to participate in top-down modulation by transmitting higher order sensory information from the LH to the AL (24, 29). However, direct evidence for top-down modulation via the CSDn has never been demonstrated. Our results suggest that during olfaction, the CSDn acts more in a bottom-up fashion where responses in the AL have a greater impact on downstream processing in the LH. Transmitter release during olfactory sampling putatively occurs only later in the sensory processing stream rather than at the earliest stages. Interestingly, the CSDn has numerous release sites in the AL (25, 30) and CSDn derived 5-HT directly inhibits several classes of neurons in the AL (30). But how are CSDn release sites activated in the AL? Olfaction clearly inhibits both the processes of the CSDn in the AL as well at its spike initiation site (30). However, it is likely that the CSDn also receives input from unidentified non-olfactory sources that excite the neuron’s spike initiation site allowing it to modulate in a top-down manner. Further reconstruction of the CSDns inputs in the EM data set will reveal candidates for further physiological evaluation. Thus, non-olfactory stimulation of the CSDn may be consistent with top-down modulation and would constitute one mechanism by which a broadly arborizing modulatory neuron may be multifunctional depending on the source of its excitatory drive. Multifunctional neurons have been well described in central pattern generating networks (68) but are only recently becoming appreciated with regards to neuromodulation (56). Local synaptic interactions within the AL could shape 5-HT release as well to alter olfactory coding in a glomerulus specific fashion (69). Our findings in *Drosophila* suggest that integration into local circuits allows modulatory cells greater flexibility in how they participate in sensory processing and may be a feature that is often overlooked when assessing the function of these critical components of the central nervous system.

## Methods

### Odors and odor delivery

Odors were presented as previously described (30). In brief, a carrier stream of carbon-filtered house air was presented at 2.2L/min to the fly continuously. A solenoid was used to redirect 200 ml/min of this air stream into an odor vial before rejoining the carrier stream, thus diluting the odor a further 10-fold prior to reaching the animal. All odors are reported as v/v dilutions in paraffin oil (J.T. Baker VWR #JTS894), except for acids, which were diluted in distilled water. All odors were obtained from Sigma Aldrich (Saint Louis, MO) except for cVA, which was obtained from Pherobank (Wageningen, Netherlands). cVA was delivered as a pure odorant. In our olfactometer design, the odor vial path was split to 16 channels each with a different odor or solvent control. Pinch valves (Clark Solutions, Hudson MA part number PS1615W24V) were used to select stimuli between each trial. Each odor was presented sequentially one trial at a time. Each odor was presented 3-4 times within a preparation and the mean of these responses were then averaged across animals.

| **Odors** | **Supplier** |
| --- | --- |
| Paraffin oil | J.T.Baker CAS: 8012-95-1 |
| Ammonium hydroxide | Sigma-Aldrich CAS: 1336-21-6 |
| Ethyl acetate | Sigma-Aldrich CAS: 141-78-6 |
| Methyl salicylate | Sigma-Aldrich CAS: 119-36-8 |
| Benzaldehyde | Sigma-Aldrich CAS: 100-52-7 |
| 3-hexanone | Sigma-Aldrich CAS: 589-38-8 |
| p-cresol | Sigma-Aldrich CAS: 106-44-5 |
| 1-octen-3-ol | Sigma-Aldrich CAS: 3391-86-4 |
| Geranyl acetate | Sigma-Aldrich CAS: 105-87-3 |
| cVA | Pherobank, Wijk bij Duurstede, Netherlands |
| geosmin | Sigma-Aldrich CAS: 16423-19-1 |
| acetic acid | Sigma-Aldrich CAS: 64-19-7 |

### Fly Genotypes

The following *Drosophila* genotypes were used in this study: w; UAS-GCaMP6s; Gal4-R60F02, UAS-GCaMP6s, w; Gal4-NP2242/Cyo; Gal4-R60F02, UAS-GCaMP6s, w; Gal4-GH146/+; UAS-GCaMP6s/+ and w; UAS-CD4:spGFP11/Q-GH146; QUAS-syb:spCFP1-10/Gal4-R60F02. To produce the singleton CSDn in Fig. 1A, we used UAS-myrGFP, QUAS-mtdTomato-3xHA/+; trans-Tango/+;Gal4-MB465C/+ which occasionally labeled individual CSDns. All flies were obtained from the Bloomington stocks repository except for NP2242 (Kyoto Stock Center).

### Calcium imaging of odor-evoked activity

Female flies aged 3-5 weeks post-eclosion and reared at room temperature were used. Imaging experiments were performed at room temperature. The brain was constantly perfused with saline containing (in mM): 103 NaCl, 3 KCl, 5 N-tris(hydroxymethyl)methyl-2-aminoethane-sulfonic acid, 8 trehalose, 10 glucose, 26 NaHCO3, 1 NaH2PO4, 2 CaCl2, and 4 MgCl2 (adjusted to 270–275 mOsm). The saline was bubbled with 95% O2/5% CO2 and reached a pH of 7.3. 920 nm wavelength light was used to excite GCaMP6s under two-photon microscopy. The microscope and data acquisition were controlled by ThorImage 3.0 (Thorlabs, Inc.). A volume of sample consisting of 6-8 frames with 5-6 μm sections was scanned at a speed of 60 frames/second. Thus, the sample was recorded at approximately 6-7 volumes/second. Single imaging trials consisted of 90 volumes at a resolution of 256×256 pixels. Odors were delivered for 1 second after the first 2 seconds of each trial. An 80 seconds of interval between each trial was applied. Calcium transients (ΔF/F) were measured as changes in fluorescence, in which ΔF/F was calculated by normalizing the fluorescence brightness changes over the baseline period (the first 2 seconds of each trial before the odor delivery). A Gaussian low-pass filter of 4×4 pixels in size was then applied to raw ΔF/F signals prior to any analysis. The frame containing the peak response was identified by plotting the averaging ΔF/F in an ROI as a function of frame number. The peak calcium signal for each trial was computed as the average of three consecutive frames centered on the frame of peak response, and we set the peak window the same for all trials within a preparation. For each odor stimulus, data was pooled by averaging the peak odor-evoked calcium signal across 3-4 repeats.

### Calcium imaging of statistical analysis

Principal Component Analysis (PCA) was applied on the spatial pattern of the peak calcium signal (ΔF/F) in each trial. PCA was computed using the “princomp” function in Matlab (Mathworks, Natick, MA). The spatial pattern was first reshaped to a one dimensional array as the input of the PCA. The output PCs were later reshaped back for display purposes. We applied Linkage Hierarchical Clustering to calculate the diversity of response patterns to different odors. The correlations between spatial response patterns (two-dimensional) were calculated as the indicator of the similarity between two odor evoked responses. The Euclidean distances between each correlation pairs were then calculated as a parameter for clustering. A regression analysis was applied to compare the similarity between the odor response patterns of the CSDn and PNs in the LH. The p-value and the square of correlation coefficient (R^2^) were calculated as the indicator of similarity for each odor response pattern pairs evoked. Two-tailed paired t-tests were performed for all comparisons between before laser ablation and after laser ablation within the same group. All statistical functions were applied in Matlab.

### CSDn EM Reconstruction

The CSDn was identified and partially reconstructed in the female adult fly brain (FAFB) dataset as described in Zheng et al 2018 using CATMAID (70, 71). The CSDn reconstruction from the cell body along the primary and secondary arbors leading into the LH as well as tertiary and quaternary branches into the lateral horn were reviewed by a second observer back to the primary branch as previously described (43). For the multi-compartmental model, measurements of the CSDn branch radius (to inform axial resistance parameters) were taken at 7 locations along the primary arbor between the contralateral AL and protocerebrum, 5 locations along the second order branch leading into the lateral horn and 6 locations along primary and secondary branches within the lateral horn. For each location, 20-50 measures of axon radius were taken from consecutive tissue sections. Data on the projection neurons that are pre-synaptic to CSDn in the FAFB dataset were provided as a personal communication from Greg Jefferis, Phillip Schlegel, Alex Bates, Marta Costa, Fiona Love and Ruari Roberts.

### Model construction

A multi-compartmental conductance-based computer model of the CSDn neuron was constructed by taking its electron micrograph reconstruction and importing it into the Neuron simulator (44). An initial reconstruction contained more than 60,000 compartments, but a simplified version of it was generated with 6,056 sections in the Neuron simulator that retained its basic anatomical and electrotonic structure. The passive cable parameters (axial resistance Ra, leak conductance gpas, leak reversal Epas, and specific capacitance Cm) were fitted using Neuron’s RunFitter algorithm. For fitting, we used recorded responses to stimuli of current-clamp and voltage-clamp steps and current-clamp white noise generated by Matlab (Mathworks, Natick, MA). Whole-cell recordings of the CSDn were performed as previously described (30). The series resistance for CSDn recordings was approximately 10MΩ, and input resistance was 500 - 600MΩ. The pipette resistance was between 8 - 10MΩ. The reversal potential of the CSDn was −45mV. While fitting, parameters were restricted by physiological ranges (R_a_ between 0.0001-5000 Ωcm, C_m_ between 0.1−2 μF/cm^2^, g_leak_ between 10^−6^−0.1 S/cm^2^, and Epas had unlimited range in mV).The resulting passive model of the CSDn neuron was simulated using Neuron’s default integration method with a time step of 0.025 ms.

To investigate how the passive signal travels from the AL to the other part of the CSDn, we randomly picked ten spots within the AL and injected a square wave of current into the model to elicit a maximum voltage response ranging from about 2 mV to 20 mV. This was repeated 6 times in each condition. We monitored the voltage changes in the windows shown in Fig. 4. The windows we set were for the cell body, the center of AL (three randomly selected monitor sites) and the LH (three randomly selected monitoring sites). The average voltage changes of each window were shown in the Fig. 4. To investigate how signals preferentially propagated along the CSDn, the same method was applied to LH, so that current was injected in the LH and voltage change windows of interests across the CSDn were kept the same. The process was then repeated with current injection into the AL.

### Laser Transection

The transection window was guided by the Gcamp6s basal fluorescence at 920 nm, at about 20 μm before the CSDn neurites enter the lateral horn. An 80mW laser pulse, which consisted of 10 repetitions of continuous frame scanning with 8 μsec of pixel dwell time, at 800 nm was then applied onto this window. A total estimated energy of 0.05 J was thus applied. Successful transection usually resulted in a small cavitation bubble (shown in Fig. 4).

### KCl induction of GRASP and Immunohistochemistry

In brief, brains used for the induction of syb:GRASP were dissected and rinsed three times with a KCl solution (42). The brains were then fixed in 4% paraformaldehyde for 20 min. We used the following primary and secondary antibodies at the indicated dilutions: 1:1000 rabbit anti-5HT Sigma (S5545), 1:50 chicken anti-GFP Invitrogen (A10262), 1:100 mouse anti-GFP (referred as anti-GRASP, Sigma #G6539, ref:3), 1:500 rat anti-N-Cadherin (Developmental Studies Hybridoma Bank, DN-Ex #8), 1:250 Alexa Fluor 633 goat anti-rabbit (Invitrogen, A21071), 1:250 Alexa Fluor 488 goat anti-chicken (Life Technologies, A11039), 1:250 Alexa Fluor 568 goat anti-mouse lgG (Life Technologies, A11004) and 1:1000 Alexa Fluor 647 donkey anti-rat IgG (AbCam, ab150155). Brains were mounted and imaged in Vectashield mounting medium (Vector Labs). All steps were performed at room temperature. Confocal z-stacks for the syb:GRASP experiments were collected with a Zeiss LSM710 microscope using a 63× oil-immersion lens and the z-stack of GFP expression in Fig. 1A was collected with an Olympus FV1000s using a 40x oil-immersion lens.

## Acknowledgement

The authors would like to thank Greg Jefferis, Phillip Schlegel, Alex Bates, Marta Costa, Fiona Love and Ruari Roberts for providing access to tracings of PNs in the FAFB dataset, as well as Jay Milam for assistance with confocal scans and Tom Kazimiers, Andrew Champions, Chris Barnes and Albert Cardona for CATMAID support and Mert Erginkaya for assistance with tracing review. This work was supported by a Whitehall Foundation Grant and an NIH R21 to QG, an NIH DC 016293 to AMD and QG, and Georgia Gwinnett College VPASA Seed Fund partially supported CG. The contributions of AMD and KEC in this project were supported in part by the Janelia Visiting Scientist Program.

---

**Figure S1.**
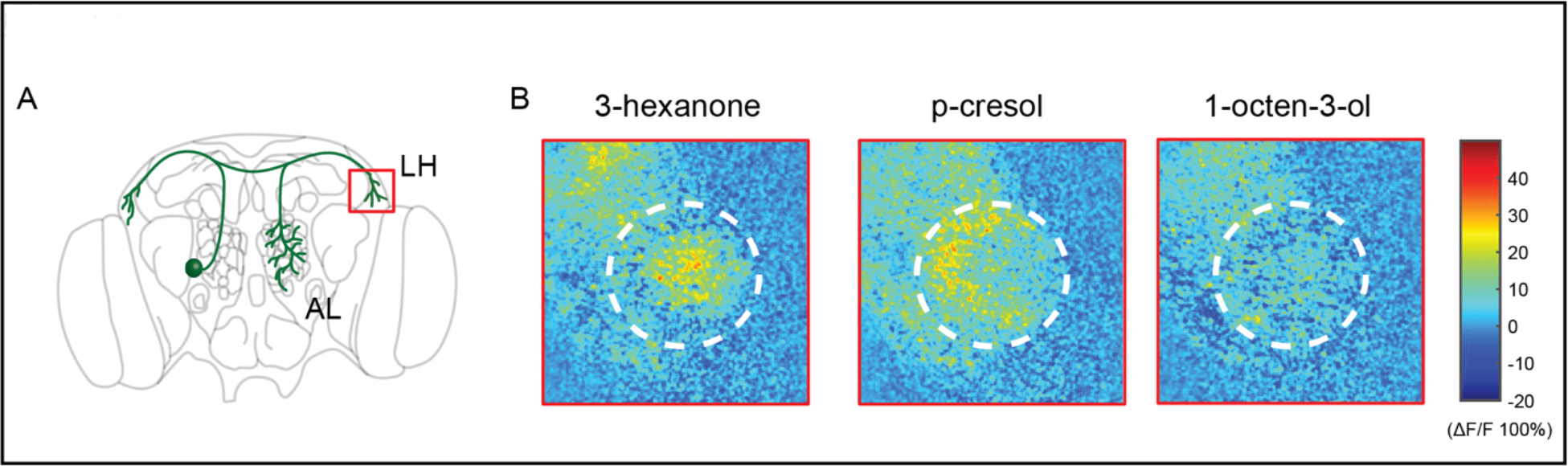
Olfactory stimulation activates presynaptic release sites in CSDn terminals in the LH. (*A*) A schematic of the *Drosophila* brain and the region of imaging. (*B*) Odor responses measured with sytGCaMP6s revealing activation of presynaptic release sites.

**Figure S2.**
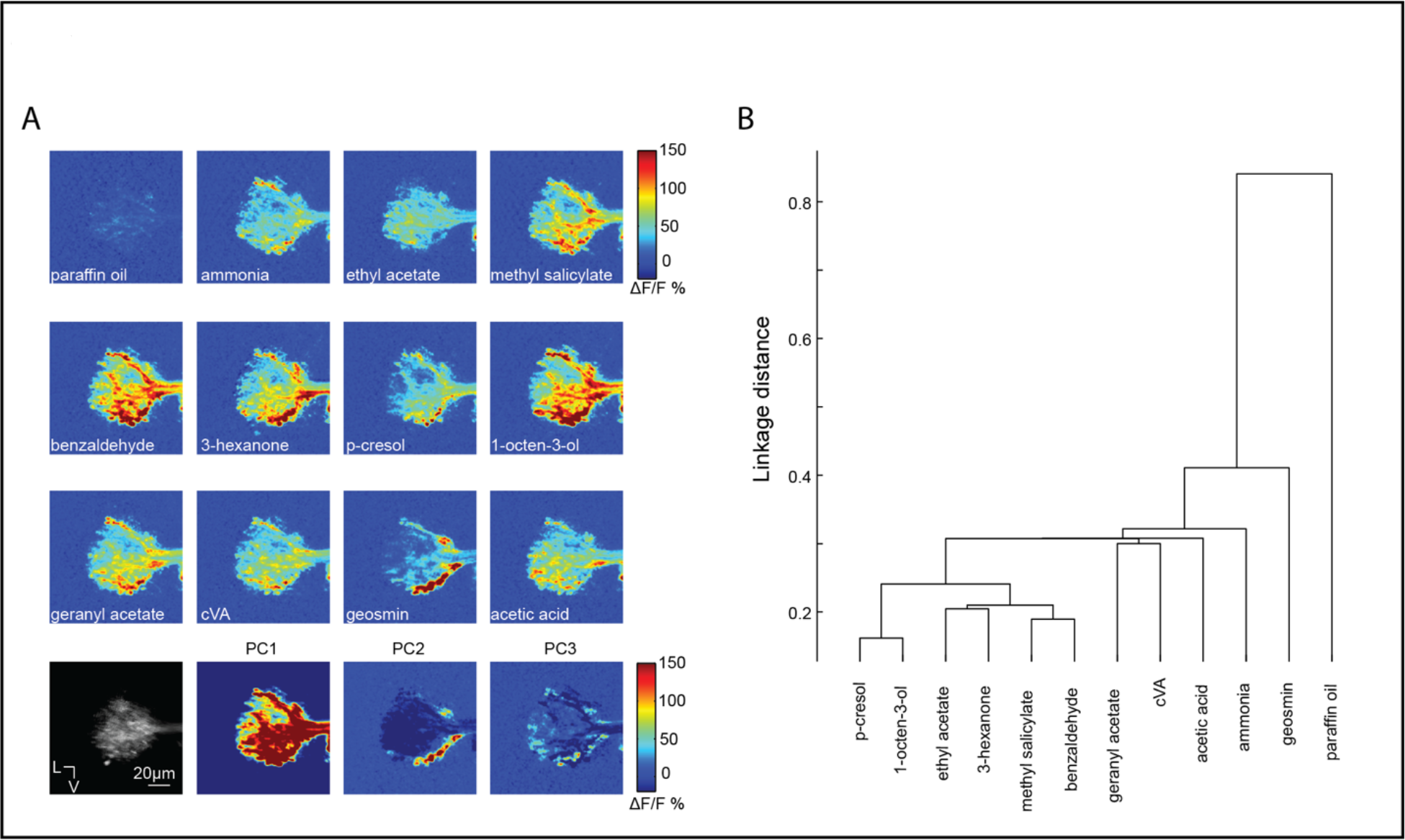
The axons of projection neurons in the LH have odor specific patterns of activation. (*A*) Calcium responses of PN axons in the LH revealed with GCaMP6s. PCA analysis shows several unique patterns of activation are required to explain the variance in the data set of odor response. (*B*) Clustering analysis on the Euclidean distance of correlations across odors in PN responses in the LH (n = 5).

**Figure S3.**
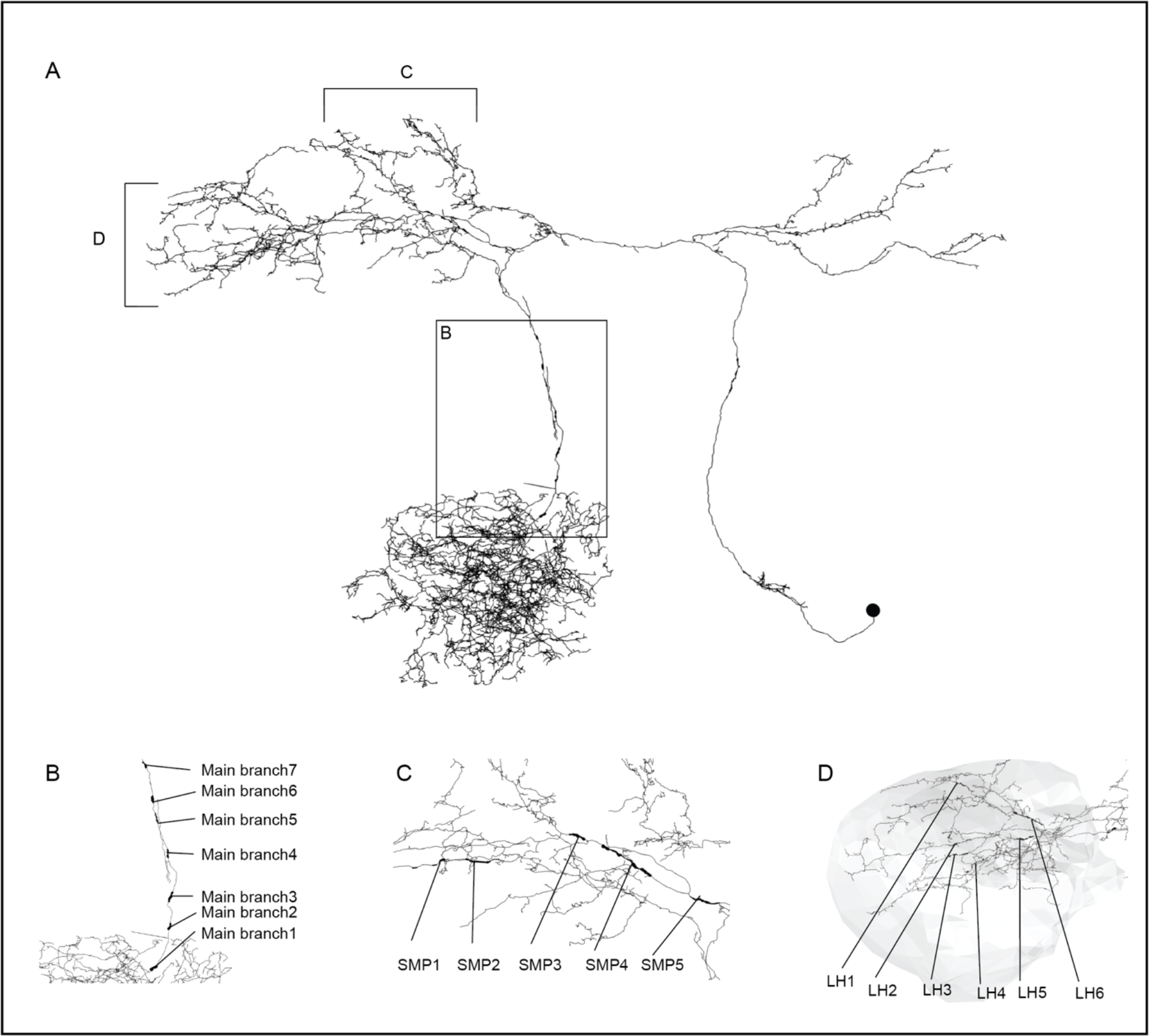
Sampling strategy for measuring CSDn axon radius. (*A*) Reconstruction of the CSDn skeleton from the whole brain EM volume. Thickened processes indicate individual sampling locations from which 20-50 measures were taken. Inset box or brackets indicate individual regions from which multiple samples were taken and shown in more detail in subsequent panels. (*B*) Axon radius sampling locations from the main branch of the CSDn. (*C*) Sampling locations from the CSDn process passing through the superior-medial protocerebrum (SMP) to the LH. (*D*) Sampling locations from 6 separate branches of the CSDn in the LH (boundary shaded in grey).

**Figure S4.**
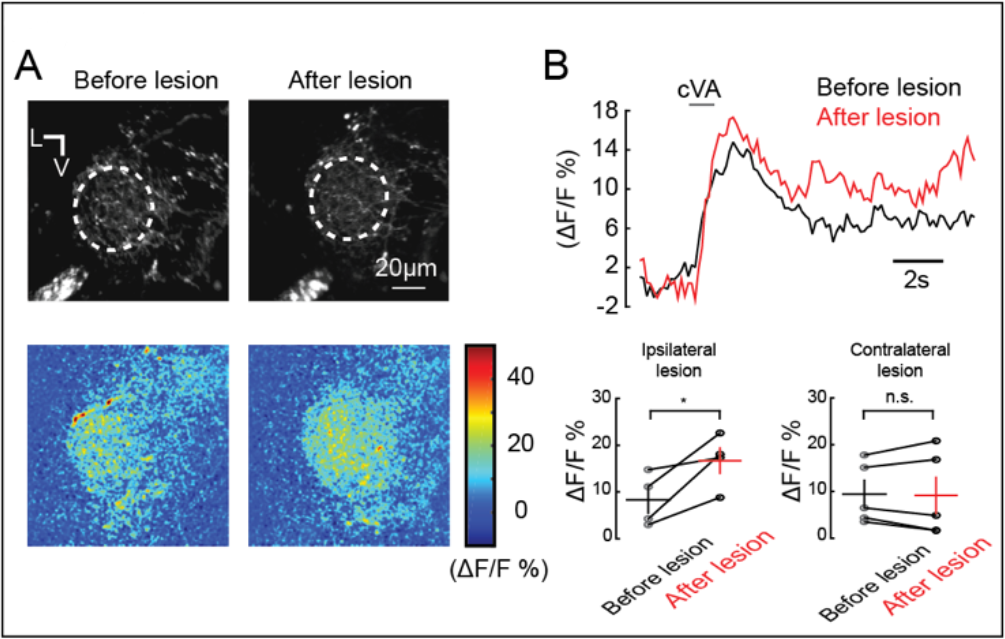
cVA excites CSDn processes in the lateral horn. (*A*) Experimental approach as in Fig. *4G*-*K*. *Top*, basal fluorescence of GCaMP6s signal before and after ablation of the processes connecting the AL and LH. *Bottom*, ΔF/F responses of CSDn neurites in the LH to cVA. (*B*) *Top*, a time course of the calcium response to cVA before and after laser ablation. *Bottom*, quantification of the data in *A* and *B* for ipsilateral and contralateral control lesion experiments.

## References

1. Katz PS (1995) Intrinsic and extrinsic neuromodulation of motor circuits. Current Opinion in Neurobiology 5(6):799–808.

2. Lizbinski KM, Dacks AM (2017) Intrinsic and Extrinsic Neuromodulation of Olfactory Processing. Front Cell Neurosci 11:424.

3. Schwarz LA, Luo L (2015) Organization of the locus coeruleus-norepinephrine system. Curr Biol 25(21):R1051–R1056.

4. Charnay Y, Léger L (2010) Brain serotonergic circuitries. Dialogues Clin Neurosci 12(4):471–487.

5. Hornung J-P (2003) The human raphe nuclei and the serotonergic system. J Chem Neuroanat 26(4):331–343.

6. Ikemoto S (2007) Dopamine reward circuitry: Two projection systems from the ventral midbrain to the nucleus accumbens–olfactory tubercle complex. Brain Research Reviews 56(1):27–78.

7. Lammel S, et al. (2008) Unique Properties of Mesoprefrontal Neurons within a Dual Mesocorticolimbic Dopamine System. Neuron 57(5):760–773.

8. Wenk GL (1997) The Nucleus Basalis Magnocellularis Cholinergic System: One Hundred Years of Progress. Neurobiology of Learning and Memory 67(2):85–95.

9. Padmanabhan K, et al. (2019) Centrifugal Inputs to the Main Olfactory Bulb Revealed Through Whole Brain Circuit-Mapping. Front Neuroanat 12:413.

10. Nunez-Parra A, Maurer RK, Krahe K, Smith RS, Araneda RC (2013) Disruption of centrifugal inhibition to olfactory bulb granule cells impairs olfactory discrimination. Proceedings of the National Academy of Sciences 110(36):14777–14782.

11. Thiele A, Bellgrove MA (2018) Neuromodulation of Attention. Neuron 97(4):769–785.

12. Jacob SN, Nienborg H (2018) Monoaminergic Neuromodulation of Sensory Processing. Front Neural Circuits 12:70–17.

13. Melzer S, et al. (2012) Long-range-projecting GABAergic neurons modulate inhibition in hippocampus and entorhinal cortex. Science 335(6075):1506–1510.

14. Miller RJ (1998) Presynaptic receptors. Annual Review of Pharmacology and Toxicology 38(1):201–227.

15. Terada S-I, Kobayashi K, Ohkura M, Nakai J, Matsuzaki M (2018) Super-wide-field two-photon imaging with a micro-optical device moving in post-objective space. Nature Communications:1–14.

16. Lecoq J, et al. (2014) Visualizing mammalian brain area interactions by dual-axis two-photon calcium imaging. Nat Neurosci 17(12):1825–1829.

17. Stirman JN, Smith IT, Kudenov MW, Smith SL (2016) Wide field-of-view, multi-region, two-photon imaging of neuronal activity in the mammalian brain. Nat Biotechnol 34(8):857–862.

18. Sofroniew NJ, Flickinger D, King J, Svoboda K (2016) A large field of view two-photon mesoscope with subcellular resolution for in vivo imaging. Elife 5:413.

19. Gagnon D, Parent M (2014) Distribution of VGLUT3 in highly collateralized axons from the rat dorsal raphe nucleus as revealed by single-neuron reconstructions. PLoS One 9(2):e87709.

20. van der Kooy D, Kuypers HG (1979) Fluorescent retrograde double labeling: axonal branching in the ascending raphe and nigral projections. Science 204(4395):873–875.

21. Otazu GH, Chae H, Davis MB, Albeanu DF (2015) Cortical Feedback Decorrelates Olfactory Bulb Output in Awake Mice. Neuron 86(6):1461–1477.

22. Rothermel M, Carey RM, Puche A, Shipley MT, Wachowiak M (2014) Cholinergic inputs from Basal forebrain add an excitatory bias to odor coding in the olfactory bulb. Journal of Neuroscience 34(13):4654–4664.

23. Dacks AM, Christensen TA, Hildebrand JG (2006) Phylogeny of a serotonin-immunoreactive neuron in the primary olfactory center of the insect brain. The Journal of Comparative Neurology 498(6):727–746.

24. Sun XJ, Tolbert LP, Hildebrand JG (1993) Ramification pattern and ultrastructural characteristics of the serotonin-immunoreactive neuron in the antennal lobe of the moth Manduca sexta: a laser scanning confocal and electron microscopic study. The Journal of Comparative Neurology 338(1):5–16.

25. Coates KE, et al. (2017) Identified Serotonergic Modulatory Neurons Have Heterogeneous Synaptic Connectivity within the Olfactory System of Drosophila. Journal of Neuroscience 37(31):7318–7331.

26. Roy B, et al. (2007) Metamorphosis of an identified serotonergic neuron in the Drosophila olfactory system. Neural Dev 2(1):1.

27. Xu L, et al. (2016) A Single Pair of Serotonergic Neurons Counteracts Serotonergic Inhibition of Ethanol Attraction in Drosophila. PLoS One 11(12):e0167518–15.

28. Singh AP, et al. (2013) Sensory neuron-derived eph regulates glomerular arbors and modulatory function of a central serotonergic neuron. PLoS Genet 9(4):e1003452.

29. Hill ES, Iwano M, Gatellier L, Kanzaki R (2002) Morphology and physiology of the serotonin-immunoreactive putative antennal lobe feedback neuron in the male silkmoth Bombyx mori. Chemical senses 27(5):475–483.

30. Zhang X, Gaudry Q (2016) Functional integration of a serotonergic neuron in the Drosophila antennal lobe. Elife 5:2435.

31. Gaudry Q (2018) Serotonergic Modulation of Olfaction in Rodents and Insects. Yale J Biol Med 91(1):23–32.

32. Chen T-W, et al. (2013) Ultrasensitive fluorescent proteins for imaging neuronal activity. Nature 499(7458):295–300.

33. Hong EJ, Wilson RI (2015) Simultaneous encoding of odors by channels with diverse sensitivity to inhibition. Neuron 85(3):573–589.

34. de Belle JS, Heisenberg M (1994) Associative odor learning in Drosophila abolished by chemical ablation of mushroom bodies. Science 263(5147):692–695.

35. Cohn R, Morantte I, Ruta V (2015) Coordinated and Compartmentalized Neuromodulation Shapes Sensory Processing in Drosophila. Cell 163(7):1742–1755.

36. Jefferis G, et al. (2007) Comprehensive maps of Drosophila higher olfactory centers: spatially segregated fruit and pheromone representation. Cell 128(6):1187–1203.

37. Tanaka NK, Awasaki T, Shimada T, Ito K (2004) Integration of Chemosensory Pathways in the Drosophila Second-Order Olfactory Centers. Current Biology 14(6):449–457.

38. Seki Y, et al. (2017) Olfactory coding from the periphery to higher brain centers in the Drosophila brain. 1–20.

39. Strutz A, et al. (2014) Decoding odor quality and intensity in the Drosophila brain. Elife 3:e04147.

40. Min S, Ai M, Shin SA, Suh GSB (2013) Dedicated olfactory neurons mediating attraction behavior to ammonia and amines in Drosophila. Proceedings of the National Academy of Sciences. doi:10.1073/pnas.1215680110.

41. Feinberg EH, et al. (2008) GFP Reconstitution Across Synaptic Partners (GRASP) Defines Cell Contacts and Synapses in Living Nervous Systems. Neuron 57(3):353–363.

42. Macpherson LJ, et al. (2015) Dynamic labelling of neural connections in multiple colours by trans-synaptic fluorescence complementation. Nature Communications 6:10024–9.

43. Zheng Z, et al. (2018) A Complete Electron Microscopy Volume of the Brain of Adult Drosophila melanogaster. Cell 174(3):730–743.e22.

44. Carnevale NT, Hines ML (2006) The NEURON Book (Cambridge University Press, Cambridge, UK).

45. Stuart G, Spruston N, Häusser M (2016) Dendrites (Oxford University Press). 3rd Ed.

46. Sparta DR, Stuber GD (2014) Cartography of Serotonergic Circuits. Neuron 83(3):513–515.

47. Pollak Dorocic I, et al. (2014) A whole-brain atlas of inputs to serotonergic neurons of the dorsal and median raphe nuclei. Neuron 83(3):663–678.

48. Weissbourd B, et al. (2014) Presynaptic Partners of Dorsal Raphe Serotonergic and GABAergic Neurons. Neuron 83(3):645–662.

49. Ogawa SK, Cohen JY, Hwang D, Uchida N, Watabe-Uchida M (2014) Organization of Monosynaptic Inputs to the Serotonin and Dopamine Neuromodulatory Systems. CellReports 8(4):1105–1118.

50. Petzold GC, Hagiwara A, Murthy VN (2009) Serotonergic modulation of odor input to the mammalian olfactory bulb. Nat Neurosci 12(6):784–791.

51. Lottem E, Lörincz ML, Mainen ZF (2016) Optogenetic Activation of Dorsal Raphe Serotonin Neurons Rapidly Inhibits Spontaneous But Not Odor-Evoked Activity in Olfactory Cortex. Journal of Neuroscience 36(1):7–18.

52. Huang Z, Thiebaud N, Fadool DA (2017) Differential serotonergic modulation across the main and accessory olfactory bulbs. J Physiol (Lond) 595(11):3515–3533.

53. Floresco SB, Yang CR, Phillips AG, Blaha CD (1998) Basolateral amygdala stimulation evokes glutamate receptor-dependent dopamine efflux in the nucleus accumbens of the anaesthetized rat. Eur J Neurosci 10(4):1241–1251.

54. Jones JL, et al. (2010) Basolateral amygdala modulates terminal dopamine release in the nucleus accumbens and conditioned responding. Biological Psychiatry 67(8):737–744.

55. Hamid AA, et al. (2015) Mesolimbic dopamine signals the value of work. Nat Neurosci 19(1):117–126.

56. Berke JD (2018) What does dopamine mean? Nat Neurosci:1–7.

57. Dankoski EC, Wightman RM (2013) Monitoring serotonin signaling on a subsecond time scale. Front Integr Neurosci 7:44.

58. Threlfell S, et al. (2004) Histamine H3 receptors inhibit serotonin release in substantia nigra pars reticulata. Journal of Neuroscience 24(40):8704–8710.

59. Schlicker E, Classen K, Göthert M (1984) Gabab Receptor-Mediated Inhibition of Serotonin Release in the Rat-Brain. Naunyn Schmiedebergs Arch Pharmacol 326(2):99–105.

60. Tao R, Auerbach SB (1995) Involvement of the dorsal raphe but not median raphe nucleus in morphine-induced increases in serotonin release in the rat forebrain. NSC 68(2):553–561.

61. Egashira N, et al. (2002) Involvement of 5-hydroxytryptamine neuronal system in Delta(9)-tetrahydrocannabinol-induced impairment of spatial memory. Eur J Pharmacol 445(3):221–229.

62. Gracia-Llanes FJ, et al. (2010) Synaptic connectivity of serotonergic axons in the olfactory glomeruli of the rat olfactory bulb. Neuroscience 169(2):770–780.

63. Suzuki Y, Kiyokage E, Sohn J, Hioki H, Toida K (2014) Structural basis for serotonergic regulation of neural circuits in the mouse olfactory bulb. J Comp Neurol 523(2):262–280.

64. Varga V, Sik A, Freund TF, Kocsis B (2002) GABA(B) receptors in the median raphe nucleus: distribution and role in the serotonergic control of hippocampal activity. NSC 109(1):119–132.

65. Ranganathan GN, et al. (2018) Active dendritic integration and mixed neocortical network representations during an adaptive sensing behavior. Nat Neurosci:1–13.

66. Jones PW, Gabbiani F (2010) Synchronized neural input shapes stimulus selectivity in a collision-detecting neuron. Curr Biol 20(22):2052–2057.

67. Koren D, Grove JCR, Wei W (2017) Cross-compartmental Modulation of Dendritic Signals for Retinal Direction Selectivity. Neuron 95(4):914–927.e4.

68. Briggman KL, Kristan WB Jr. (2008) Multifunctional pattern-generating circuits. Annu Rev Neurosci 31:271–294.

69. Kloppenburg P, Mercer AR (2008) Serotonin modulation of moth central olfactory neurons. Annu Rev Entomol 53(1):179–190.

70. Schneider-Mizell CM, et al. (2016) Quantitative neuroanatomy for connectomics in Drosophila. Elife 5. doi:10.7554/eLife.12059.

71. Saalfeld S, Cardona A, Hartenstein V, Tomancak P (2009) CATMAID: collaborative annotation toolkit for massive amounts of image data. Bioinformatics 25(15):1984–1986.

